# A high-throughput, image-based assay to assess drug sensitivity of *Acanthamoeba castellanii* cysts

**DOI:** 10.1101/2025.06.06.658337

**Authors:** Carrie A. Flynn, Andrew Harmez, Rebecca I. Colón-Ríos, William F. Flynn, Barbara I. Kazmierczak

## Abstract

Pathogenic free-living amoebae (FLA) such as *Acanthamoeba castellanii* are present in soil and water worldwide. *A. castellanii* causes systemic infections with very high mortality rates, yet drugs specifically targeting this pathogen are not available. Methods to reliably generate and assay cysts, which drive infection recurrence and drug resistance, are unavailable in a high-throughput format suitable for drug screening and testing. In this study, we developed a robust and reproducible protocol for encysting *A. castellanii* as well as a high-throughput, quantitative cyst viability assay using fluorescent live/dead staining coupled with microscopy and automated image analysis. These methods were used to screen the cysticidal activity of 17 clinically relevant drugs and disinfectants and identified five agents, including caspofungin, as active against cysts.

## Background

The pathogenic free-living amoeba (FLA) *Acanthamoeba castellanii* causes disseminated infections, granulomatous amebic encephalitis (GAE), and contact lens-associated keratitis. With a mortality rate >90% for systemic infections despite multidrug therapy, active drugs against *A. castellanii* are urgently needed (1, 2). Drug discovery requires quantitative, high-throughput assays to reliably measure viability of the three *A. castellanii* life forms involved in infection: vegetative trophozoites, inactive pseudocysts, and dormant cysts. Cysts are particularly difficult to kill and cause recurrent infections (3, 4). We recently developed a method to measure live vs. dead trophozoites, live vs. dead pseudocysts, and live cysts(5). Although this method avoids falsely scoring viable cysts as “dead”, it fails to accurately score cyst killing. To date, therefore, no reliable and quantitative viability assay allows high-throughput drug screening on cysts.

Cyst physiology is a major barrier to measuring cyst death. When motile and metabolically active trophozoites encounter environmental stress they differentiate into either pseudocysts (6) or cysts. Differentiation can be triggered by changes in pH or temperature, drug exposure, or starvation. Trophozoites shed large amounts of water, develop a double cell wall, and firmly surface-attach as they differentiate into metabolically inactive cysts with a cellulose-rich inner cell wall (the endocyst) and a protein-rich outer cell wall (the ectocyst). These changes help cysts resist amoebicides, both environmental and pharmaceutical. Dormant cysts can survive for years awaiting favorable environmental conditions, which cause them to excyst and return to the trophozoite form (7–9).

The lack of robust viability assays and standardized methods for inducing uniform and complete trophozoite encystment have made studies of cysts challenging (10–12). Existing protocols often lead to incomplete encystment (with immature cysts lacking the signature double cell wall), asynchronous encystment (leading to a mix of trophozoites, immature cysts, and mature cysts), and/or excessive cell death. Established techniques for assessing cyst viability such as plaque assays(13) and cell counting with trypan blue are low-throughput, labor intensive, and often unreliable (14, 15) while assays that measure metabolic activity falsely score dormant cysts as “dead”. The proteinaceous ectocyst wall remains attached to surfaces after cyst death, causing assays that measure cell adherence (e.g. SRB) to report dead cysts as live(5). Variation among methods used to induce encystment and quantify cyst viability has made it difficult to compare and reproduce data between labs.

We set out to develop a robust and synchronous encystment protocol, as well as a quantitative cyst viability assay suitable for high-throughput drug screening. Our method uses an image segmentation tool(16) that performs automated image analysis of fluorescently stained cysts. This method was validated and then used to evaluate the cysticidal activity of 17 clinically relevant drugs and disinfectants.

## Materials and Methods

### Amoeba culture

Amoebas were cultured as described in (5). Encystment was induced in sterile, flat-bottom, tissue culture-treated 96-well plates (clear walls/bottoms for SRB; clear bottoms/black walls for fluorescent staining). Trophozoites were harvested and plated as previously described (5) at a density of 10^5^ cells/well, using trophozoites from a single mother flask to encourage synchronous encystment. Plated trophozoites were centrifuged at 200 xg for 5 minutes and incubated at room temperature for 2 hours to allow cell attachment. PYG medium was then removed and replaced with 150 µL per well FEM unless otherwise indicated. Media was always added gently to well walls to avoid detaching adhered cells. Plates were incubated with breathable plate seals (Axygen BF-400) instead of lids for 72 hours at 25^◦^C. Plates were then used for experiments as described below.

In experiments involving SDS treatment to lyse immature cysts and trophozoites, cysts were induced in plates as described above. At the end of the 72-hour encystment, media was removed and half the wells per condition were incubated with 150 µL per well 0.5% (w/v) SDS in sterile dH_2_O. Plates were incubated at room temperature on a gyratory platform for 10 minutes. SDS was removed and excess was washed away by submerging plates in PBS-MC twice before fixing with 10% TCA; the SRB assay was then performed to quantify remaining adherent cells.

To generate autoclaved negative controls, cysts induced in flasks in FEM for at least 72 hours were harvested as above, using PBS for washing and harvesting steps because FEM components precipitate when autoclaved. Cysts were autoclaved in 10-mL glass screw-top vials on a liquid cycle for 15 minutes, media volume lost to boiling was replaced, and 100 µL per well was added to 96-well plates.

### Compound testing

FEM was aspirated from encysted cells, and 100 µL per well PBS (with or without drugs) was gently added to well walls. Autoclaved cysts (control) were added to plates at this time to avoid pelleting or loss of dead cells. Plates were incubated for 20 hours at 25^◦^C without shaking, with breathable plate seals instead of lids. Spent media was removed and discarded, and cyst viability assayed by fluorescent live/dead staining protocol.

### SRB Assay

The SRB assay was performed as described in (5). A detailed protocol is provided in supplemental materials.

### Cyst viability assay

Following incubation of cysts with media or drugs, media was aspirated and wells washed once with 150 µL of sterile PBS; this minimized background fluorescence from residual compounds/medium. Invitrogen LIVE/DEAD staining reagent was prepared by mixing 4 µM ethidium homodimer-1 (EthD-1) and 2 µM calcein-AM (c-AM) in sterile PBS. Wells were stained with 50 µL per well LIVE/DEAD reagent and incubated in the dark for 40-60 minutes at room temperature. Phase-brightfield and fluorescent (using GFP and TRITC channels to image c-AM and EthD-1 staining, respectively) images of wells were then taken with an automated Nikon Eclipse Ti-E inverted microscope (20x objective) and an Andor Zyla VSC-01400 camera with 3 separate Z stacks taken with 5 µm steps per channel, per well. Exposure times were 50 ms for phase and GFP images and 500 ms for TRITC images. Images were then analyzed as described below.

### Image segmentation and quantification

Images were acquired on a Nikon Ti PFS-S (0.45 NA, Andor Zyla VSC-01400 camera) microscope with a 20x objective (Plan Fluor ELWD 20x DIC L) as described above. Each image series contains between 1 and 96 images, corresponding to wells in 96 well plates, and each series image contains between 1 and 5 z planes, one brightfield phase-contrast (PC) channel, and two color channels (TRITC red and GFP green).

Multi-series Nikon .nd2 files were converted to single series 3-channel OME TIFF files using BioFormats bfconvert(17) and condensed to a single z-plane using maximum intensity projection. The PC channel of each image was transformed using CLAHE adaptive histogram equalization and normalized by its maximum intensity, then used as input to cyto2 model of CellPose(16) without default percentile normalization using an approximate object diameter size of 30 pixels. The resulting segmentation PC mask was used to mask the two color channels, allowing tabulation of per-object and background quantification metrics. Only objects with equivalent area diameter between 10 and 25 µm were included for downstream quantification. In plates with mean background intensity below 1000FU, all objects with mean red fluorescence intensity greater than 2500FU were classified as stained in order to account for a large dynamic range of staining intensity across dead cysts. Segmented objects in plates with either highly variable background intensity or background intensity greater than 5000FU were classified as stained using a per-well threshold of 1500FU above the mean background intensity. These parameters were tuned over a training set of 188 images and validated against manual counts of 57 images containing 12,444 cells. This pipeline was then used to evaluate a set of 2794 images containing approximately 2.3 million cells. Downstream collation, quantification, and analysis was performed using the Python scientific programming stack(18).

### Calcofluor white staining

Encysted cells were removed by scraping, pelleted, resuspended in PBS, and stained with 20 µM calcofluor white (Biotium) per manufacturer’s instructions. Briefly, cells were incubated with calcofluor white for 20 min at room temperature, with rocking. 2 µL of cells were spotted onto a 1.5% low-melt agarose pad (AmericanBio) (19), inverted onto a MatTek 35 mm uncoated glass-bottom dish (no. 1.5 coverslip, 20 mm diameter), and imaged immediately using a Nikon Eclipse Ti-E inverted microscope (100x objective with oil) and a Andor Zyla VSC-01400 camera using the phase and DAPI channels.

### Data analysis

Data were graphed and analyzed using GraphPad Prism version 10. Mean and standard error of the mean (SEM) are displayed for each scatterplot. Statistical significance was determined for two groups using a two-tailed, unpaired t test with Welch’s correction and for three or more groups using one-way Brown-Forsythe and Welch ANOVA tests (for unequal SDs) with either Dunnett’s T3 (for n<50 per group) or Games-Howell (for n>50 per group) comparisons tests; an alpha of 0.05 was used to determine significance.

### Code availability

The code used to extract, segment, quantify, and collate results is available at https://github.com/wflynny/amoeba-segmenter.

### Data availability

Tabular per-object segmentation properties and quantification measurements for all experiments is available at https://github.com/wflynny/cyst-viability-assay.

## Results

### Systematic optimization of an encystment protocol

Our goal was to develop a 96 well plate-based high-throughput assay for cyst viability. As *Acanthamoeba* adheres tightly to surfaces as it encysts, but cysts cannot reattach once removed(20) (Fig S1A), we developed a method to encyst cells in microtiter plates. Aspects of 96-well cultivation (e.g. low oxygen, variable oxygen between wells, high density) hinder synchronous cyst induction and no protocols have described reliable *A. castellanii* encystment in microtiter plates.

We used an EMb media-based microplate encystment protocol (5) as a starting point for optimization. Encystment efficiency was measured by SRB assay ± 0.5% SDS; this detergent selectively lyses trophozoites and immature cysts(21) (Fig S3). We saw significant well-to-well variation with EMb, with trophozoites, immature cysts, and mature cysts all present in non-SDS treated wells at 48h (Fig S2D). Next, substrates and cofactors known to upregulate the major enzymes involved in cyst wall construction, e.g. cellulose synthase (8, 9), glycogen phosphorylase(22), sirtuins(23), and several proteases(24, 25) were systematically added to EMb encystment media (Table S1). Factors that upregulate cellulose synthase in plants were tested(26–28), along with sugar and amino acid building blocks for components of the outer cyst wall (12). Encystment triggers, e.g. increased osmolarity, stress, and low oxygenation, were likewise varied (22, 29). We used an iterative process that quickly removed failing conditions, while ingredients that promoted encystment were further combined and tested, for a total of 835 media combinations. SRB assay results were confirmed by visualization of the characteristic cyst double cell wall and wrinkled, star-shaped endocyst in all cells.

45 media formulations (Table S2) had high absorbance (≥2.0) by SRB assay for all replicate wells at 48-72h of incubation (Fig S4), with four media leading to mean absorbance values ≥2.5 at 72 hours of incubation (Fig 1A). These four media were each applied to a quadrant of a 96-well plate to assess heterogeneity by plate position (Fig 1B). One formulation (#33) performed significantly better than the others and was adopted for use in all subsequent experiments (“Flynn’s encystment medium” (FEM)). Microscopy demonstrated a monolayer of mature cysts for FEM (#33) vs. mixtures of cysts and trophozoites for media #8, 11, and 25 (Fig 1C). In these latter media trophozoites grazed on cysts, underscoring how asynchronous encystment provided food (cysts) that prevented trophozoite encystment. Encystation of all cells required 72 hours of incubation in FEM (Fig 1D) and yielded cells with the classic morphology of a mature cyst and bright calcofluor white staining (Fig 1E). This method therefore achieved the synchronous induction of mature, viable cysts in microtiter plates with significantly improved cyst survival and adherence over previous methods (Fig 1F).

**Figure 1:**
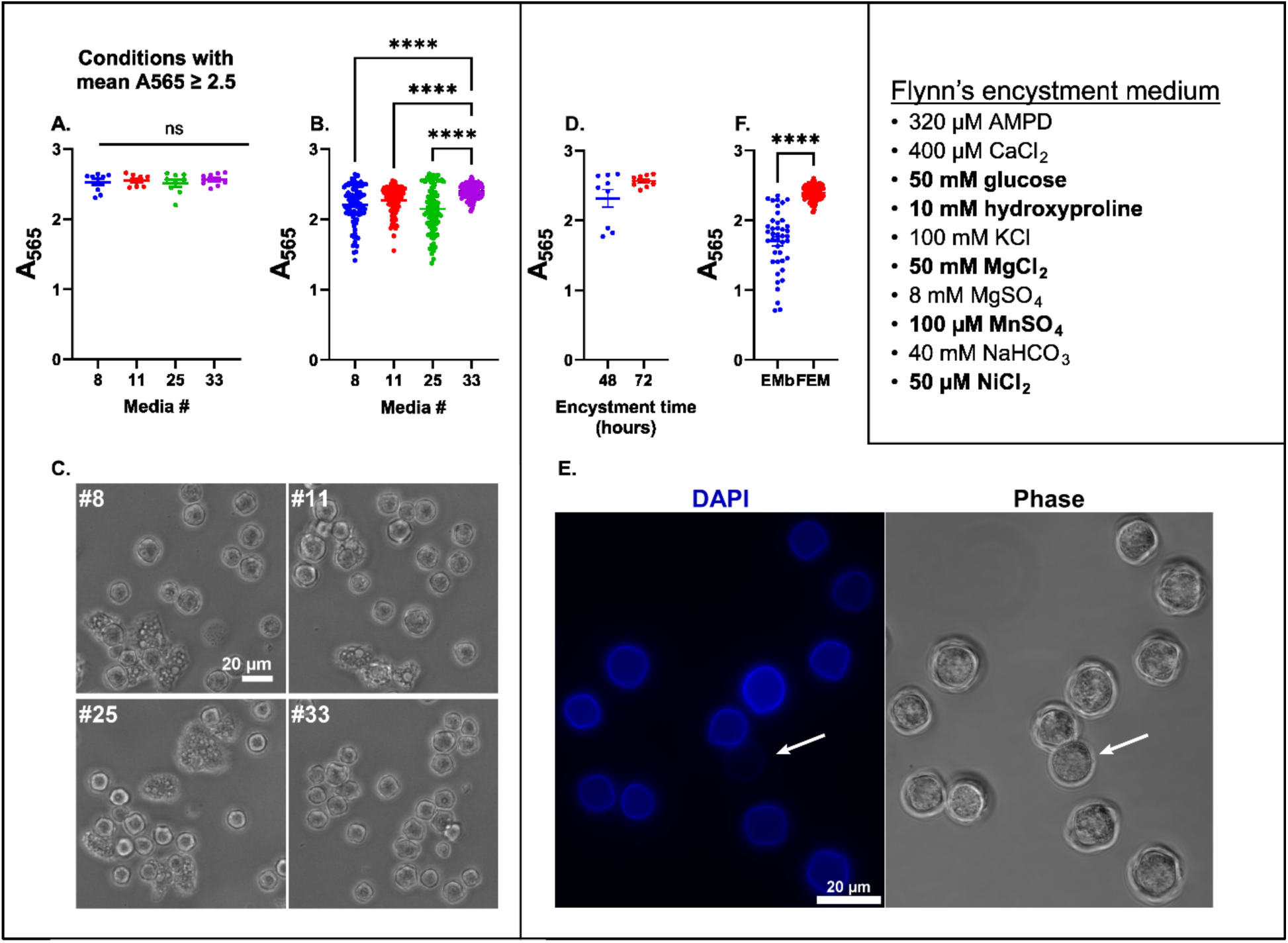
FEM induces synchronous, complete encystment. **A:** Conditions from figure S4 with a mean A565 ≥ 2.5. Components of each formulation listed in Table S2. **B:** Trophozoites were encysted in the same 4 conditions shown in A, but with 24 replicate wells per condition to assess for variability in plate location. **C:** Microscopy of cells from B at end of encystment. Phase images acquired using an EVOS FL Auto Imaging System (20x objective). **D:** Cells were encysted in FEM (ingredients listed in inset; bolded items are new components unique to FEM, unbolded items are ingredients in EMb) for 48 (blue) or 72 (red) hours. **E:** Cells encysted in FEM were stained with 20 µM calcofluor white and imaged using a Nikon Eclipse Ti-E inverted microscope (100x objective with oil) in a glass-bottom dish overlaid with a 1.5% agarose pad. Representative images displayed. Arrow, immature cyst. **F:** Cells were encysted using our former protocol in EMb for 48 hours (blue) or our optimized method in FEM for 72 hours (red). **A, B, D, F:** Scatter plots report the absorbance measured by SRB assay after SDS treatment. The mean and SEM of at least 3 independent experiments with at least 3 replicate wells per condition are shown. Statistical differences between 3 or more groups calculated with one-way ANOVA (Brown-Forsythe and Welch ANOVA tests for unequal SDs) with Dunnett’s T3 multiple comparisons test (A) or with Games-Howell multiple comparisons test (B, for n>50); differences between 2 groups calculated with two-tailed, unpaired t test with Welch’s correction (F); ns for not significant (p>0.05), **** for p<0.0001.

### Optimization of a viability assay

Our next goal was to develop a quantitative cyst viability assay suitable for high-throughput drug screening in microtiter plates. We first evaluated fluorescent live/dead staining with calcein-AM (c-AM) and ethidium homodimer-1 (EthD-1), as this method has been successfully used with a range of cell types(30). EthD-1 uniformly stained autoclaved *A. castellanii* cysts and rarely stained viable cysts (Fig S5), as expected. Calcein-AM, however, failed to stain live cysts; the few cells that were c-AM-positive usually also stained with EthD-1. We hypothesized that c-AM did not penetrate an intact double cyst wall, but could enter cysts that had compromised cell walls because they were dying or excysting.

Images of stained cysts that had been incubated in FEM or in varying concentrations of the cysticidal agents chlorhexidine or H_2_O_2_ were manually counted, yielding numbers of total (Fig S6A) and red-stained cells (Fig S6B). We then calculated the percent of cysts stained red by EthD-1 (Fig S6C). These manually annotated images were used to train a model to discriminate live and dead cells (Fig S6C). There was excellent correlation between our manual counts and those generated by the automated pipeline (Fig 2A), with a Spearman Rho statistic of 0.917 (p<0.0001) and a Pearson R statistic of 0.985 (p<0.0001), as calculated on percent red cells.

**Figure 2:**
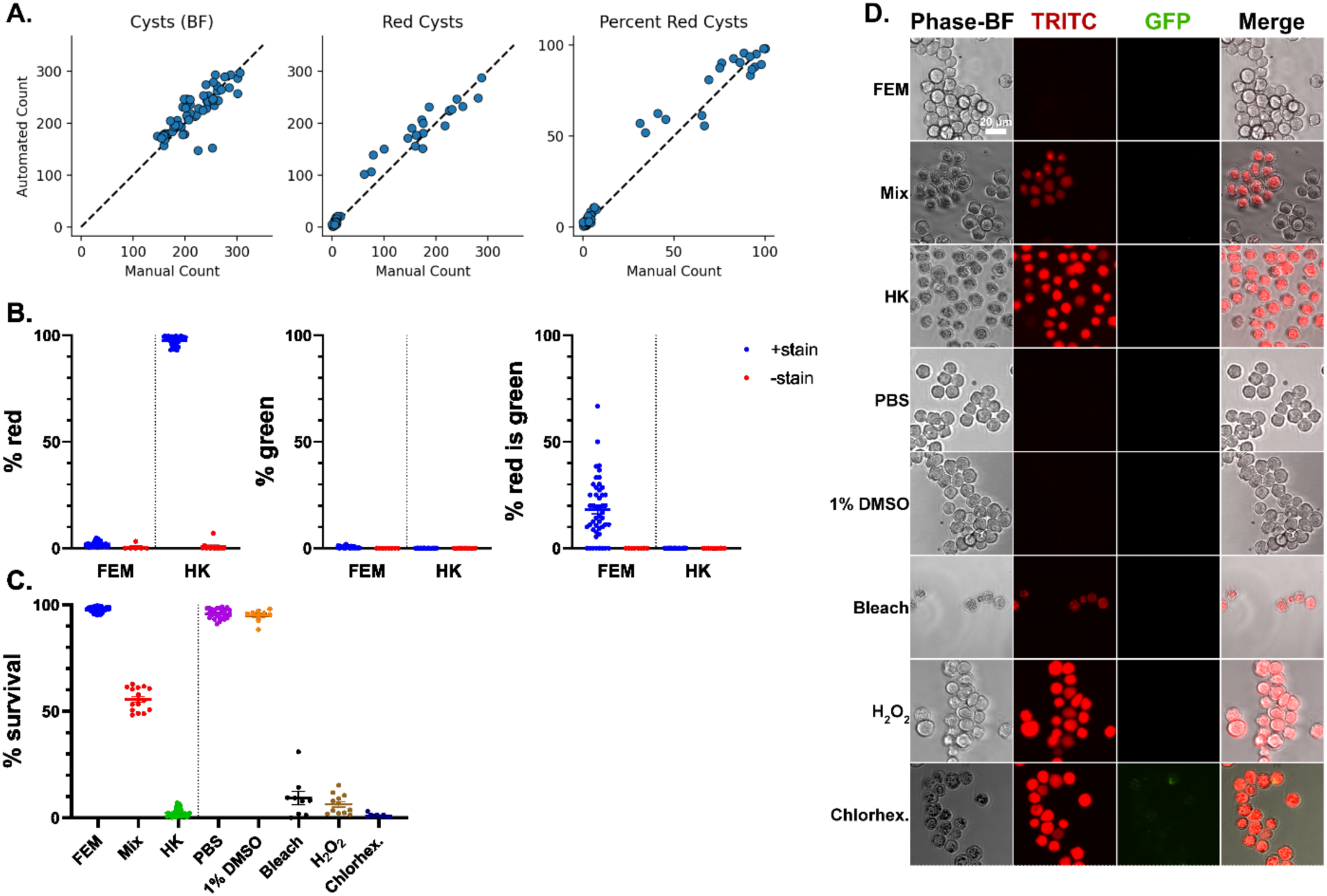
EthD-1 staining and automate d image analysis accurately measure dead cysts. **A:** The manual cell counts (x axis) reported in figure S6 were compared to those generated using the automated image analysis pipeline (y axis). Dashed line represents the perfect fit line y = x. **B:** Quantification of cysts described in figure S5. Live cysts in FEM and those that were heat-killed (HK) by autoclaving were incubated with fluorescent live/dead stain (blue) or PBS control (red), imaged with a Nikon Eclipse Ti-E inverted microscope (20x objective), and counted using automated image analysis pipeline. The mean and SEM of at least 3 independent experiments with at least 3 replicates per condition are shown. % red, percent of total cells stained with EthD-1; % green, percent of total cells stained with c-AM; % red is green, percent of red cells that also stained with c-AM. **C, D:** Cysts were treated with 0.5% bleach, 10% H_2_O_2_, 20 µM chlorhexidine, or media controls for 20 hours and compared to live cysts in FEM, heat-killed (HK) cysts, or a 1:1 mix of HK and live cysts in FEM (mix). Cells were imaged and stained with the fluorescent live/dead staining assay and counted using automated image analysis pipeline. Chlorhexidine was dissolved in DMSO; bleach and H_2_O_2_ were dissolved in PBS; all three were added at 1% final volume in PBS. **C:** Percent survival calculated by subtracting the percent of EthD-1-stained cells (measured by fluorescent live/dead staining) from 100%. The mean and SEM of at least 3 independent experiments with at least 3 replicates per condition are shown. Statistical differences calculated with one-way ANOVA (Brown-Forsythe and Welch ANOVA tests for unequal SDs) with Dunnett’s T3 multiple comparisons test. **D:** Microscopy of cysts following treatment. Images acquired using a Nikon Eclipse Ti-E inverted microscope (20x objective). Representative images displayed.

The image analysis pipeline was applied to test images (Figure S5) and confirmed our manual assessment (Fig 2B). 97.6% of autoclaved cysts, but only 2% of cysts in FEM, stained red with EthD-1 (Fig 2B, % red). There was little background fluorescence, with <1% of unstained controls scored as red in either group. Only 0.45% of live cysts in FEM and no autoclaved cells or unstained controls were scored as green (Fig 2B, % green). As we observed, a sizable fraction (18.1%) of red-stained cells were dually labeled with c-AM (Fig 2B).

The live/dead staining assay and automated image analysis pipeline were subsequently used to measure cyst viability in different media conditions, following exposure to various cysticides, and for samples constructed from live and dead cysts (Fig 2C). The method performed well, reporting high survival for media control wells, low survival (determined by percent of cells stained red) for cells killed by autoclaving (heat-killed, HK), bleach, chlorhexidine, and H_2_O_2_, and intermediate survival for mixtures of live and dead cells. Bleach (0.5%) appeared to dissolve cysts, with treated wells showing few cell bodies that stained dimly with EthD-1 (Fig 2D), likely due to nucleic acid degradation. This assay thus relies on the presence of intact dead cell bodies. Imaging again demonstrated the lack of c-AM staining of live or dead cysts.

### Outgrowth experiments confirm the specificity of EthD-1 for dead cells

We compared the results of our EthD-1 based viability assay with outgrowth experiments over a range of conditions. Live cysts and dead (autoclaved) controls were tested along with cysts treated with drugs or chemical disinfectants. Microscopic images underwent both automated computational pipeline analysis as well as manual inspection to assess cell morphology and its correlation with fluorescent staining. In parallel, both spent medium (containing detached cells) and adherent cells/material were assayed for growth in rich medium over a 2-week period following chemical or drug treatment, allowing us to evaluate regrowth of all cells originally present in an experimental well.

The EthD-1 based viability assay reported high survival for control wells containing cysts in FEM, PBS, 1% DMSO, or PYG, and low survival for autoclaved cysts (Fig 3, scatter plot). Phase-brightfield microscopy revealed a monolayer of live cysts in FEM control wells, while trophozoites and excysting cells were present in PYG wells (Fig S7B). Autoclaved cysts appeared granular and deformed; they stained with EthD-1 but not c-AM (Fig S7A, B). Treatment with bleach, PHMB, chlorhexidine, or H_2_O_2_ efficiently killed cysts according to the viability assay, and these cells appeared dead by phase imaging. While most disinfectant-treated cells stained exclusively with EthD-1, some cysts incubated in these conditions showed both EthD-1 and c-AM staining (non-representative images selected to demonstrate phenotype in Fig S7B, quantified in Fig S7A). These may represent dying cells that have not yet lost esterase activity. A high background c-AM signal was seen in wells containing H_2_O_2_, which we ascribed to peroxide-induced cleavage of c-AM. LNS-AmB and fluconazole exhibited no cysticidal activity, while caspofungin and miltefosine both resulted in partial killing, which was confirmed by microscopy.

**Figure 3:**
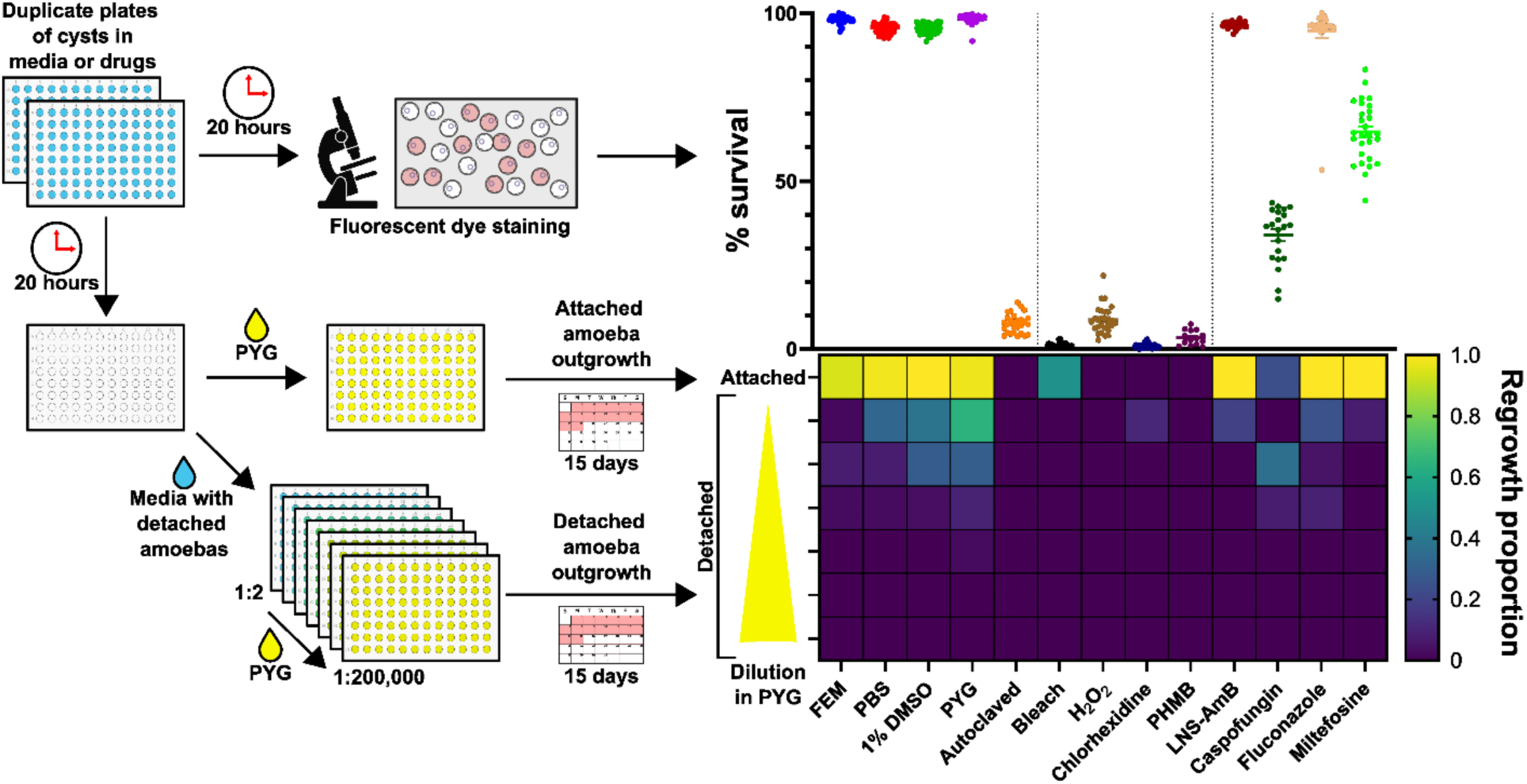
Fluorescent live/dead staining accurately labels dead cysts. Schematic of experiment illustrates experimental design and corresponding results. 10^5^ cysts per well were incubated in duplicate 96-well plates for 20 hours in conditions as indicated. For one plate, spent media was removed and the fluorescent live/dead staining assay was performed (scatter plot). For the second plate, the top 50 µL of spent media was transferred to a new plate and the remainder was removed and discarded. The transferred media was mixed 1:1 with fresh PYG, and serially diluted. Fresh PYG was also added back to wells of the original plate (attached). Plates were incubated and monitored for growth for 15 days, with fresh PYG added daily to feed cells and maintain volume. The proportion of wells that regrew during this time was scored for each condition (heat map). Amphotericin B (lipid nanosphere formulation, “LNS-AmB”), fluconazole, miltefosine, chlorhexidine, and caspofungin were dissolved in DMSO; bleach, PHMB, and H_2_O_2_ were dissolved in PBS; all were added at 1% final volume in PBS at the following concentrations: 0.25% bleach, 10% H_2_O_2_, 20 µM chlorhexidine, 50 ppm PHMB, 256 µM LNS-AmB, 64 µM caspofungin, 512 µM fluconazole, 256 µM miltefosine. **Scatter plot:** Percent survival calculated by subtracting the percent of EthD-1-stained cells (measured by fluorescent live/dead staining) from 100%. Data expressed as the percent of cells surviving at the end of incubation for each experiment; the mean and SEM of at least 3 independent experiments with 7 replicates per condition are shown. **Heat map:** Outgrowth of viable amoebas. Heatmap shows the proportion of wells that regrew during the 15-day incubation.

Cells incubated in PYG, FEM, PBS, and 1% DMSO demonstrated regrowth of trophozoites in both the original plate and in replated media samples, while no outgrowth was observed for wells containing autoclaved cysts (Fig 3, heat map). No growth in the original plate or of replated media at any dilution was detected for PHMB or H_2_O_2_, confirming the viability assay results (Fig 3, scatter plot). Regrowth of trophozoites was observed for all other conditions, indicating the presence of live cells at the end of incubation, either attached (if outgrowth occurred in the original plate) or floating (if outgrowth occurred in the removed/replated media diluted with PYG).

Both LNS-AmB and fluconazole treatment resulted in outgrowth of attached and detached cells (Fig 3, heat map), consistent with the lack of killing as measured by our viability assay (Fig 3, scatter plot). Outgrowth was also observed for wells treated with either miltefosine or caspofungin, consistent with partial killing measured by EthD-1 staining. Three of the 168 wells containing detached cells and none from the original plate treated with chlorhexidine resulted in regrowth of trophozoites. This is consistent with effective killing by chlorhexidine as reported by EthD-1 staining and illustrates the ability of outgrowth experiments to detect low numbers of viable cells at the end of drug treatment.

Many wells treated with bleach showed trophozoite outgrowth despite low survival per the EthD-1 assay. Bleach was tested at its estimated MIC (0.5%), as higher concentrations dissolved cyst cells and interfered with staining. Manual inspection of microscopy images showed that bleach-treated cells that stained with both EthD-1 and c-AM were brighter in the GFP channel and more weakly fluorescent in the TRITC channel than dually stained cells exposed to chlorhexidine, PHMB, or H_2_O_2_ (Fig S7B). Similarly, dually-stained cells treated with caspofungin (which killed only a fraction of cysts at this concentration) also exhibited brighter c-AM fluorescence. We hypothesize that dually labeled cells exhibiting bright green fluorescence were dying but still viable, while those with weak green fluorescence were no longer viable. Miltefosine-treated cells showed another staining pattern, with no c-AM staining of either EthD-1-positive or -negative cells. This variation in c-AM staining, along with the high GFP-channel background fluorescence in wells containing many dead cells, complicated the interpretation of green-stained cells. We therefore only used the proportion of EthD-1-stained cysts to quantify cell death in subsequent compound testing. c-AM staining was noted and used to flag samples where additional testing might be required to distinguish dying versus dead cells.

### Evaluating the cysticidal activity of clinically relevant drugs

Seventeen clinically relevant drugs were evaluated for cysticidal activity. Chemical disinfectants as well as commercially available antibiotic, antiparasitic, and antifungal agents with potential activity against *A. castellanii* were tested over a range of concentrations, allowing an IC50 to be calculated for each drug that killed at least 50% of the cells (Fig 4). IC50s for cysts were compared to those previously reported for trophozoites (Table 1)(5). Live versus dead cell morphology was confirmed by manual inspection of microscopy images (Fig S8).

**Figure 4:**
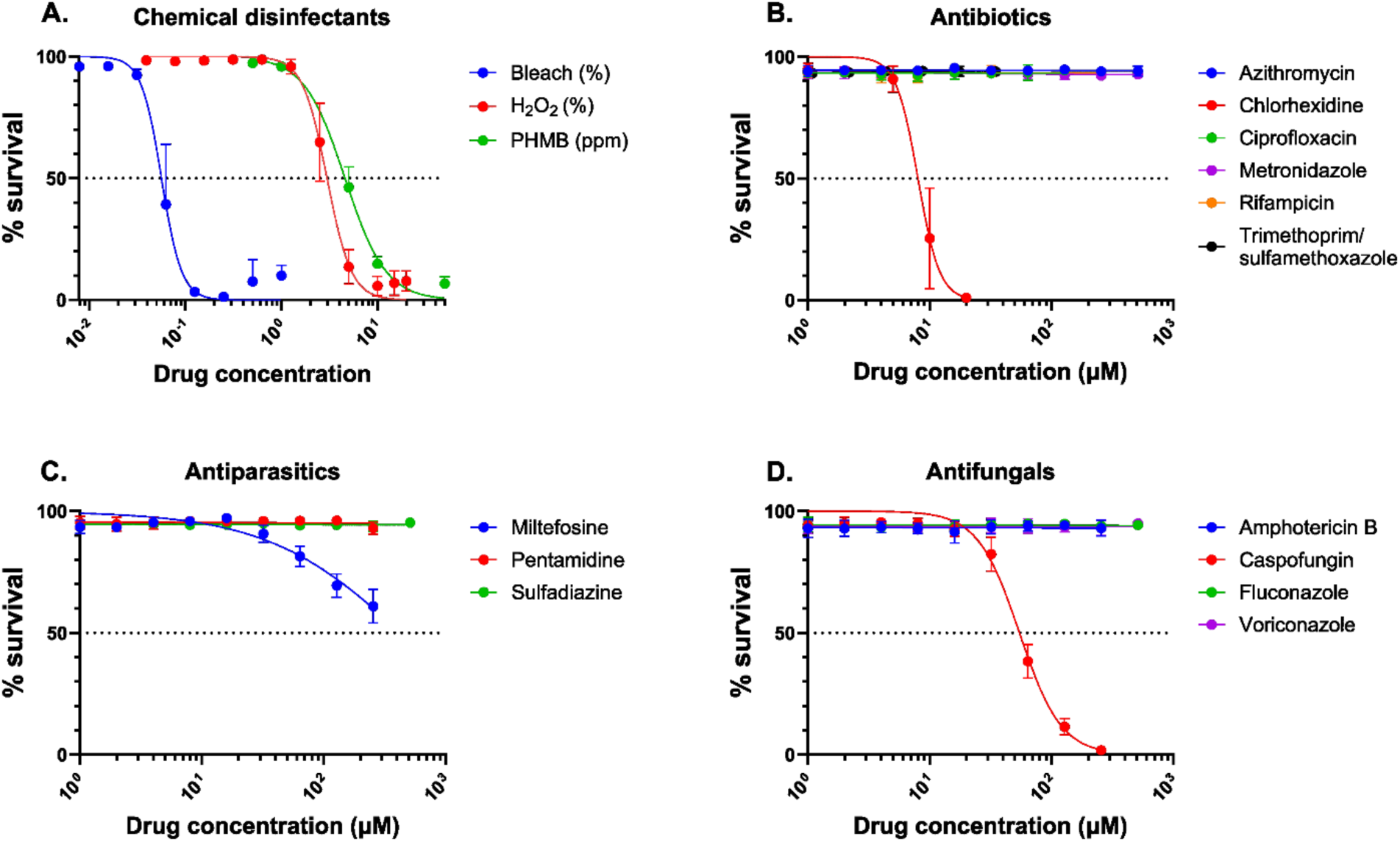
Assessing the cysticidal activity of drugs. Cysts were incubated in 96-well plates for 20 hours at 25^◦^C in conditions as indicated. At the end of the incubation, spent media was removed and the fluorescent live/dead staining assay was performed. Fluconazole, voriconazole, miltefosine, chlorhexidine, sulfadiazine, azithromycin, ciprofloxacin, metronidazole, rifampicin, trimethoprim/sulfamethoxazole, and caspofungin were dissolved in DMSO; bleach, H_2_O_2_, pentamidine, and PHMB were dissolved in PBS; amphotericin B (deoxycholate formulation) was purchased as a solution in water. Two-fold dilutions were prepared, and drugs were added at 1% final volume in PBS at a range of concentrations (listed in Table 1) according to their solubility and activity. Trimethoprim was combined with sulfamethoxazole at a ratio of 1:5.7; concentration plotted is of trimethoprim. Data expressed as the percent of cells surviving at the end of incubation as measured by fluorescent live/dead staining. The mean and SD of at least 3 independent experiments with at least 3 replicates per condition are shown.

**Table 1:** IC50 of tested compounds on cysts and trophozoites.

| Drug | Concentration range tested (cysts) | IC50 cysts | IC50 trophozoites |
| --- | --- | --- | --- |
| Amphotericin B (deoxycholate) | 1-256 μM | ND | ND |
| Azithromycin | 1-512 μM | ND | ND |
| Bleach | 0.0078125-1% | 0.05626% | 0.3277% |
| Caspofungin | 1-256 μM | 54.61 μM | 42.31 μM |
| Chlorhexidine | 1-20 μM | 8.019 μM | 4.29 μM |
| Ciprofloxacin | 1-128 μM | ND | ND |
| Fluconazole | 1-512 μM | ND | ND |
| Hydrogen peroxide | 0.039-20% | 2.984% | 0.004494% |
| Metronidazole | 1-512 μM | ND | ND |
| Miltefosine | 1-256 μM | ND | 44.99 μM |
| Pentamidine | 1-256 μM | ND | 23.03 μM |
| PHMB | 0.5 ppm (0.00005%)-600 ppm (0.06%) | 0.0004588% (4.588 ppm) | NA |
| Rifampicin | 1-256 μM | ND | ND |
| Sulfadiazine | 1-512 μM | ND | ND |
| Trimethoprim/<br>sulfamethoxazole<br>(TMP/SMX) | 1.09375/6.25-35/200 μM | ND | ND |
| Voriconazole | 1-512 μM | ND | ND |

IC50 of each compound tested on cysts, calculated by nonlinear regression using least squares fit. Previously published IC50s for trophozoites(5) are listed for comparison; drugs with activity
on both life forms are highlighted. ND, not determined (maximum concentration tested did not result in death of at least 50% of cells); NA, not applicable (drug not tested on trophozoites).

All three chemical disinfectants (bleach, H_2_O_2_, and PHMB) efficiently killed cysts (Fig 4). The concentration of H_2_O_2_ required to kill cysts was significantly higher than that for trophozoites (Table 1); surprisingly, bleach killed cysts at a lower concentration than trophozoites. Only two other agents exhibited cysticidal activity, i.e. chlorhexidine and caspofungin, at higher concentrations than those required to kill trophozoites. All other tested drugs failed to kill > 50% of the cysts after 20 hours, including the trophocidal agents pentamidine and miltefosine(5).

Wells treated with the highest concentration of amphotericin B (deoxycholate) contained rare trophozoites (Fig S8), which was not seen in any other condition. Overall, EthD-1-staining was strongly correlated with dead cell morphology, while c-AM staining varied among different treatments. At concentrations closest to the IC50, caspofungin-, chlorhexidine-, and PHMB-treated samples showed many dually-stained cysts; this was also seen for some cells treated with H_2_O_2_. Wells with the fewest green-labeled cells had the most green background fluorescence. This may reflect different mechanisms of cell death: dying cells that retain an intact envelope might also retain cytoplasmic contents (including esterases) and dually fluoresce, while lysing cells could release still-active esterases into the culture medium, leading to decreased c-AM staining and high green background fluorescence.

Overall, only five of the 17 drugs and chemical disinfectants tested demonstrated cysticidal activity and only one (chlorhexidine) was highly active with an IC50 less than 10 μM. Only one systemic agent, caspofungin, had activity against both trophozoites and cysts.

## Discussion

We have developed a highly reproducible protocol to encyst *A. castellanii* trophozoites in a format amenable for high-throughput experimentation and coupled it with a quantitative, high-throughput assay for cyst viability. These innovations support both the development of critically needed pharmaceuticals with cysticidal activity as well as broader studies of cyst biology.

We systematically approached the challenge of developing a reliable encystment protocol, optimizing environmental variables and rationally modifying media composition to promote encystment. Our best-performing encystment medium, FEM, included compounds specifically targeting biosynthetic pathways required for encystment. Low concentrations of NiCl_2_ (50 μM) and MnSO_4_ (100 μM) enhanced the activity of AcLAP, a leucine aminopeptidase involved in encystment (25). MgCl_2_, reported to trigger encystment (likely through its role as a cofactor for proteases), was included (21, 24, 29) as was MnSO_4_, a cofactor for highly expressed cellulose synthase (27). FEM also contained hydroxyproline, a significant component of the ectocyst(12). As autophagy is activated during encystment and several proteases are highly upregulated, perhaps to provide sufficient amino acids to build the outer cyst wall(31, 32), we provided these building blocks in many of our tested encystment media. Casamino acids were an ingredient in several of the best-performing media we tested, and almost all contained either casamino acids or hydroxyproline likely because of their contribution to ectocyst formation. Glucose, which has been reported to stimulate encystment when provided in the context of starvation signals (21), may help build the cellulose-rich inner cyst wall. High glucose also increases osmolarity, which independently promotes encystation(33). While we also tested many sugars that are minor components of the endocyst, we ultimately found that glucose alone performed best.

An important aspect of our work was the development and validation of a high-throughput method and analytic pipeline to identify viable but metabolically inactive cyst cells. Our assay exploited the strong attachment of encysted cells to a surface (even after cell death) and the differential permeability of live vs. dead cysts to EthD-1. We found that this nucleic acid stain was both a sensitive and specific marker of cyst death. While mammalian cell studies commonly couple EthD-1 staining with flow cytometry, this approach is technically challenging for highly adherent cysts, especially if assayed in a high-throughput 96-well format. We therefore turned to image-based quantification of stained cells.

Cysts lack a nucleus, have a visible double cell wall with a wrinkled or star-shaped inner cell wall, are small and rounded, and encyst in clusters. Standard pipelines for cell-based image analysis such as CellProfiler (ilastik) and ImageJ performed poorly when applied to these non-mammalian cells. We therefore developed a custom pipeline using CellPose for this task, which was highly scalable, tunable (segmentation and quantification are separated), and relatively performant. These features made it resilient to diverse conditions, such as high proportions of dead cells, excessive precipitated material, and high cell density. Although this analysis pipeline could not identify trophozoites, due to their variable cell size and shape, it—coupled with our highly efficient encystment protocol—made high-throughput screening for cysticidal compounds feasible.

We used our method to screen 17 clinically relevant drugs and disinfectants for cysticidal activity. Only 5 of these agents were active, including one systemic drug (caspofungin) and one compound with an IC50 less than 10 μM (chlorhexidine). Our measured IC50 for caspofungin (54.61 μM or 59.7 ng/mL) may(34) or may not(35) be achievable in the CNS, highlighting the need for development of novel therapeutics. To our knowledge, this clinically available antifungal has not been used to treat a patient with an *Acanthamoeba* infection, although our findings of both trophocidal and cysticidal activity for caspofungin may prompt consideration of its use, especially as published screens have not identified compounds active against both life forms(36). We also observed that treatment with amphotericin B (deoxycholate) uniquely resulted in the presence of trophozoites at the end of incubation. This drug might provide a nutritional or other environmental cue to excyst and further argues against its use in treating *Acanthamoeba* infections(1, 2).

In summary, we have now developed a reproducible, synchronous encystment protocol and robust method to measure viability of *Acanthamoeba* cysts. Coupled with prior methods to measure trophozoites and pseudocysts(5), this work provides essential tools that support drug development for an incurable infection.

