## Supplemental Materials for "A high-throughput, image-based assay to assess drug sensitivity of *Acanthamoeba castellanii* cysts"

Supplemental Material for “A high-throughput, image-based assay to assess drug sensitivity of *Acanthamoeba castellanii* cysts”

Carrie A. Flynn, Andrew Harmez, Rebecca I. Colón-Ríos, William F. Flynn, Barbara I. Kazmierczak

Supplemental Figures

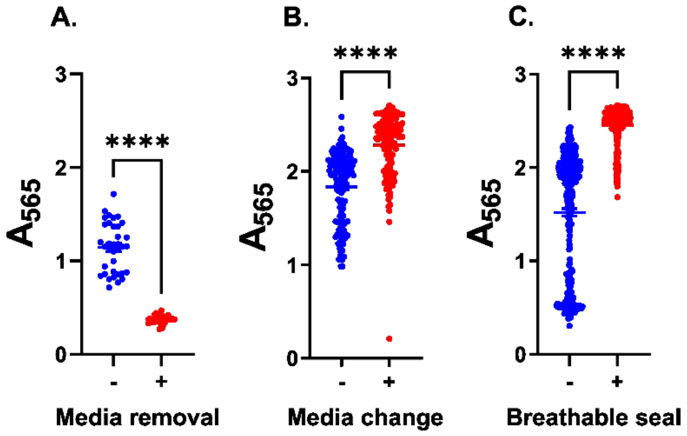


**Figure S1: In situ encystment with even oxygenation is necessary for cell adherence. A:** Trophozoites were encysted for at least 3 days in a T75 growth flask in EMb medium, removed by cell scraping, and transferred to 96-well plates. Cells were centrifuged to the bottoms of the wells and incubated at 25°C for 24 hours to allow time for cell attachment. Cells were then fixed by adding TCA to the wells with (red) or without (blue) removing medium first. **B:** Harvested trophozoites were resuspended in EM medium, added to plates, and centrifuged to well bottoms (blue) or were plated in PYG growth medium and centrifuged to well bottoms before medium was aspirated and replaced with EM medium. Cells were encysted at 25°C for 30 hours before fixing with TCA. **C:** Plated trophozoites were encysted in EM medium for 24 hours at 25°C with either the plate lid (blue) or a breathable plate seal (red) before fixing with TCA. Scatter plots report the absorbance measured by SRB assay. The mean and SEM of at least 3 independent experiments with at least 12 replicate wells per condition are shown. For B and C, half of a plate was used for each condition to assess variability by plate location, which remains high. Statistical differences calculated with two-tailed, unpaired t test with Welch's correction using GraphPad Prism software; **** for $p<0.0001$.

Cysts were induced in growth flasks by removing the spent PYG from a confluent monolayer of trophozoites, washing once with EMb, and adding 40 mL fresh EMb. Flasks were incubated without shaking at 25°C for at least 3 days until all trophozoites had encysted (determined by microscopic examination). Cysts were washed once, and 20 mL fresh EMb was added to flasks before harvesting cysts by cell scraping. Cells were counted with a hemocytometer and trypan blue dye, pelleted by centrifuging at 1,500 xg for 20 minutes at room temperature, and adjusted to a concentration of $10^6$ cysts per mL in fresh EMb. 100 μL per well was added to sterile tissue culture-treated 96-well plates with clear walls and flat bottoms for a final concentration of $10^5$ cells per well. Plates were spun at 200 xg for 5 minutes (Thermo Scientific Sorvall ST 40R Centrifuge) and immediately fixed either by adding 25 μL 50% TCA in PBS-MC per well directly to medium (blue) or by removing the culture medium and adding 125 μL 10% TCA in PBS-MC (red); the SRB assay was then performed.

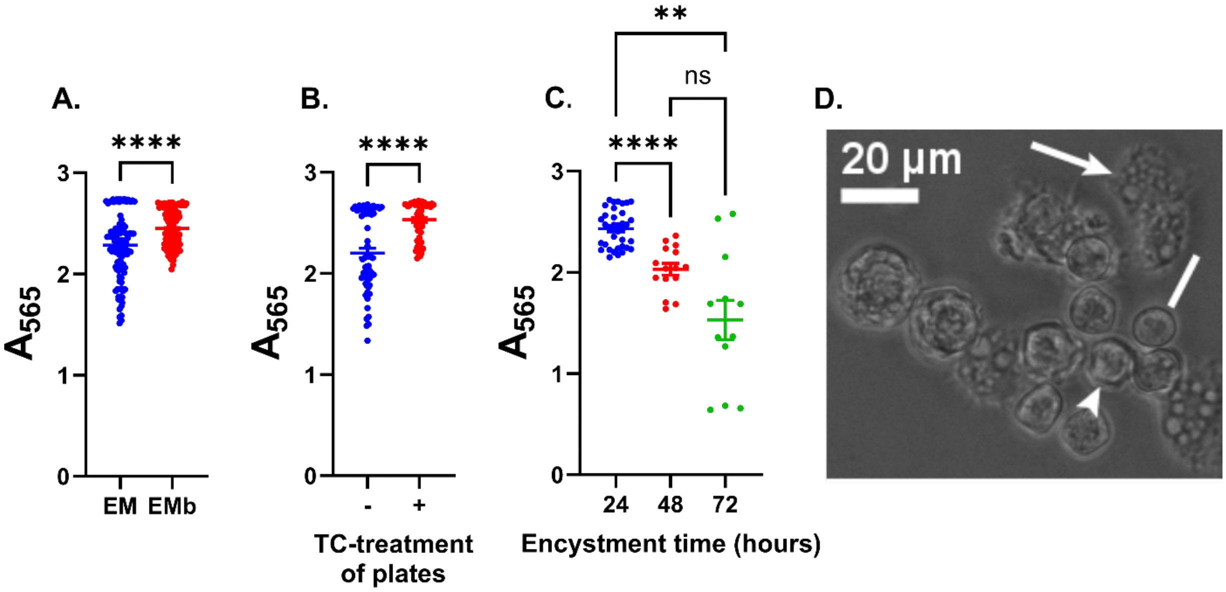


**Figure S2: Further optimization improves cell adherence but does not lead to complete encystment. A:** Plated trophozoites were encysted in EM, buffered by Tris base (blue), or EMb, buffered by AMPD (red), medium for 24 hours at 25°C before fixing with TCA. **B:** Trophozoites were dispensed in tissue culture-treated (red) or untreated (blue) plates, encysted in EMb medium for 24 hours, and fixed with TCA. **C:** Trophozoites were encysted in EMb medium according to previously optimized conditions for 24 (blue), 48 (red), or 72 (green) hours and fixed with TCA. **D:** Microscopy of cells encysted for 48 hours according to optimized protocol. Phase image acquired before spent media was removed from wells using an EVOS FL Auto Imaging System (20x objective). Arrow, trophozoite; arrowhead, mature cyst; line, immature cyst. Scatter plots report the absorbance measured by SRB assay. The mean and SEM of at least 4 independent experiments with at least 3 replicate wells per condition are shown. Statistical differences calculated with two-tailed, unpaired t test with Welch's correction (A and B) or with one-way ANOVA (Brown-Forsythe and Welch ANOVA tests for unequal SDs) with Dunnett's T3 multiple comparisons test (C) using GraphPad Prism software; ns for not significant ($p>0.05$), ** for $p<0.01$, **** for $p<0.0001$.

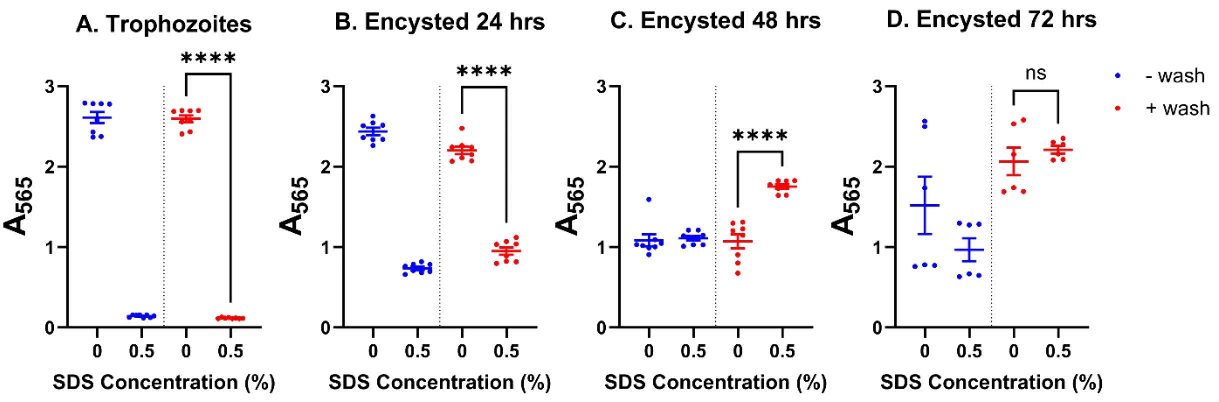


**Figure S3: Mature cysts resist SDS treatment.** Trophozoites (A) or cells encysted in EMb medium according to previously optimized conditions for 24 (B), 48 (C), or 72 (D) hours at 25°C were treated with 0.5% SDS for 10 minutes. Half the plates were washed twice by submerging in PBS-MC (red) before fixing with TCA, SDS was removed from the other half without washing before fixation (blue). Scatter plot reports the absorbance measured by SRB assay. The mean and SEM of 2 independent experiments with at least 3 replicate wells per condition are shown. Statistical differences calculated with two-tailed, unpaired t test with Welch's correction using GraphPad Prism software; **** for $p<0.0001$, ns for not significant.

Trophozoites treated with SDS (A, red) all lysed, as demonstrated by the low A565 values (confirmed by microscopic inspection). Cells encysted for 24 hours (B, red) consisted of a mix of trophozoites, immature cysts, and mature cysts, reflected by the significant drop in A565 upon treatment with SDS (lysis of trophozoites and immature cysts) while still having a higher absorbance value than SDS-treated trophozoites (A, red). The A565 of SDS-treated cells continues to increase as the number of mature cysts rises from 48 (C, red) to 72 (D, red) hours of encystment. By 72 hours of encystment, there is no significant difference between SDS-treated and untreated cells, reflecting complete encystation. The increase in absorbance values between untreated and SDS-treated cells at 48 hours is somewhat surprising and may be due to the removal of salts that accumulate as well volume decreases with evaporation over time (and may interfere with SRB dye binding). The lower values seen for unwashed (blue) vs. washed (red) wells following SDS treatment across all timepoints demonstrates interference with SRB dye binding by SDS.

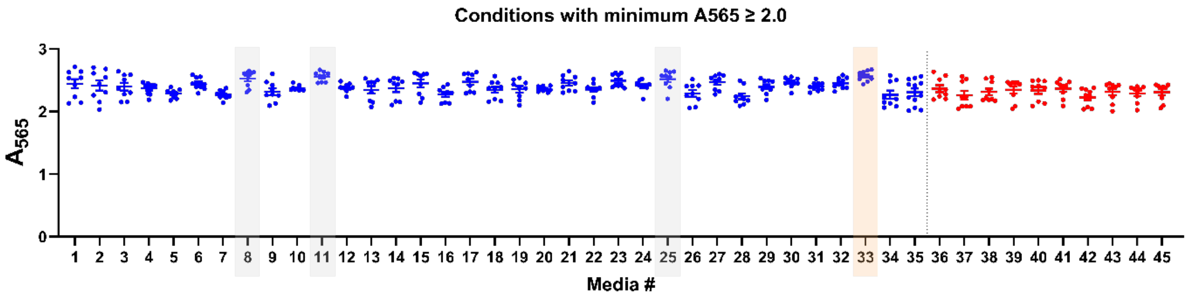


**Figure S4: Best-performing encystment media candidates.** Trophozoites were incubated in encystment media for 72 (#1-35, blue) or 48 (#36-45, red) hours at 25°C, treated with 0.5% SDS for 10 minutes (controls without SDS not shown), washed twice by submerging in PBS-MC, and fixed with TCA. Scatter plot reports the absorbance measured by SRB assay for all conditions with a minimum A565 ≥ 2.0. The mean and SEM of 3 independent experiments with 3 replicate wells per condition are shown. Bars highlight the 4 conditions with mean A565 ≥ 2.5; peach highlights the final formulation FEM (discussed in figure 1). Components of each formulation listed in Table S2.

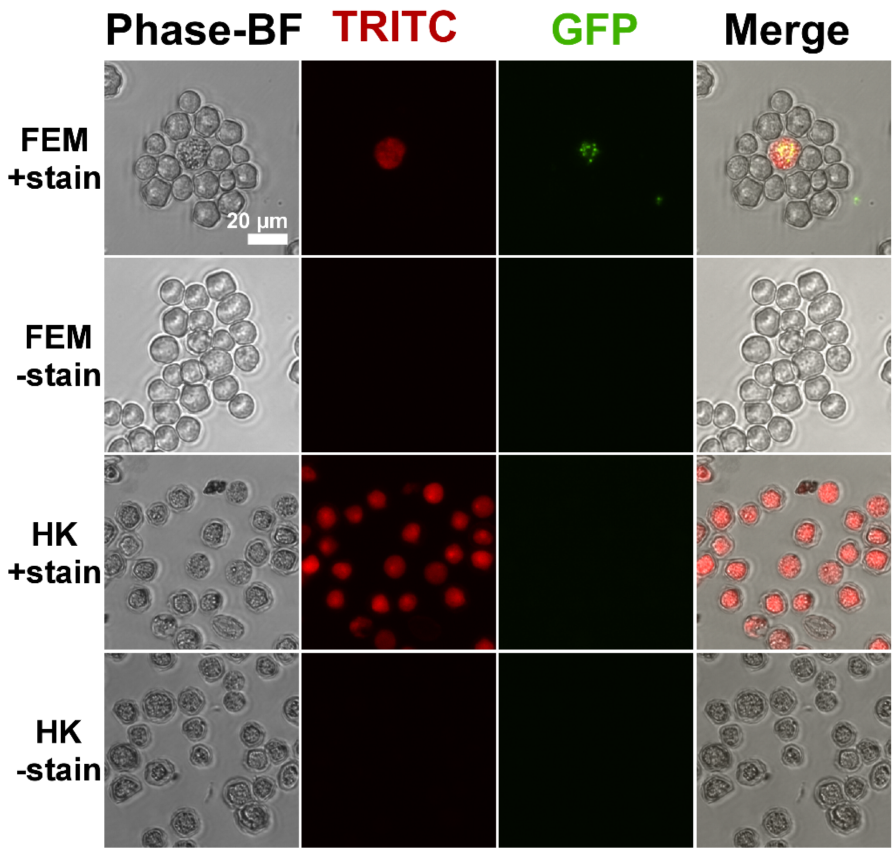


**Figure S5: EthD-1 selectively stains dead cysts.** Live cysts in FEM and those that were heat-killed (HK) by autoclaving were incubated with fluorescent live/dead (+stain) or PBS control (-stain) and imaged with a Nikon Eclipse Ti-E inverted microscope (20x objective) using the phrase-brightfield, TRITC (for EthD-1 staining), and GFP (for c-AM staining) channels. Representative images displayed.

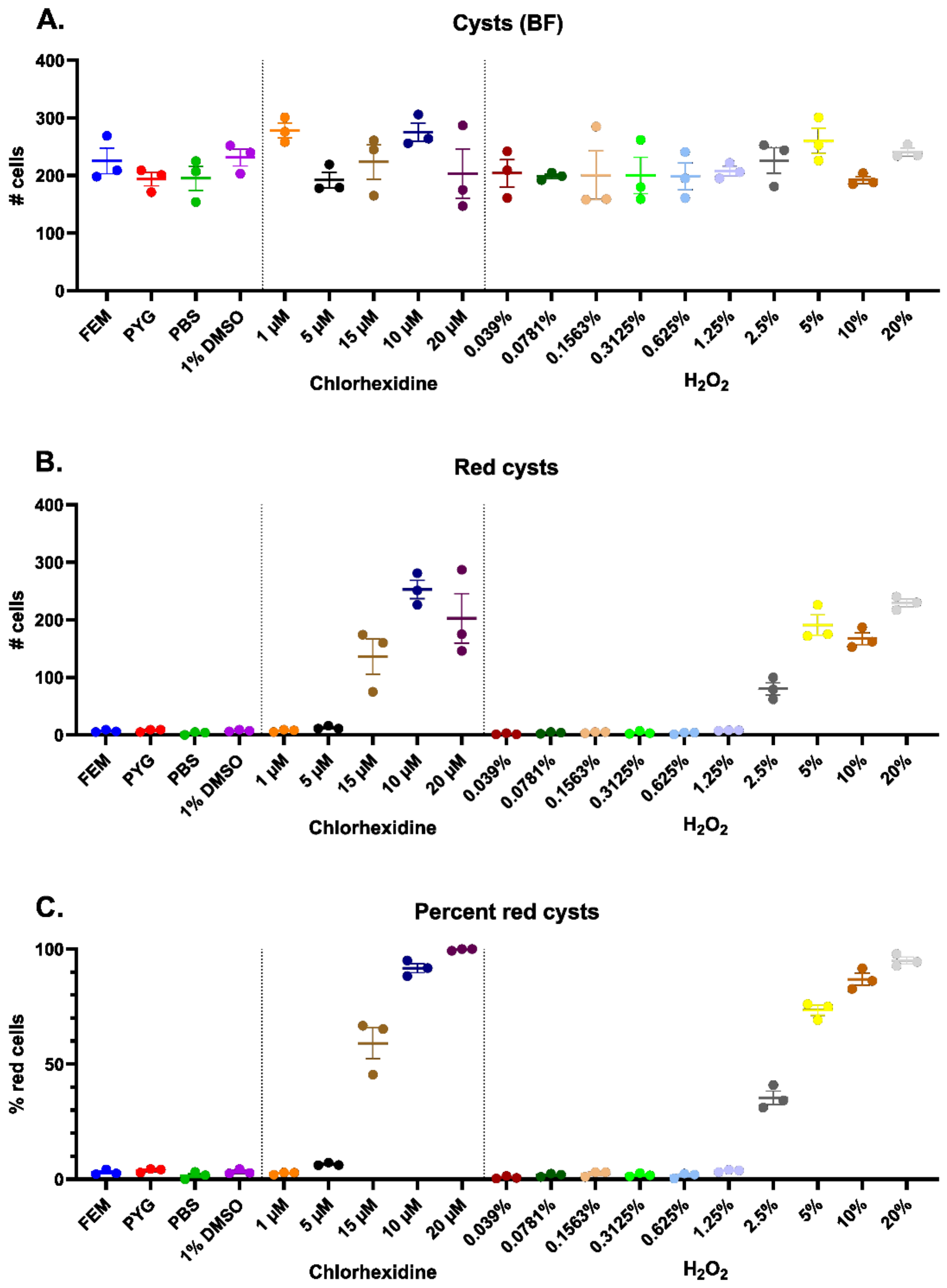


**Figure S6: Manual counts of cyst images.** Cysts were incubated for 20 hours in conditions as indicated, fluorescent live/dead stained, and imaged with a Nikon Eclipse Ti-E inverted microscope (20x objective). Images were divided into quadrants and one quadrant from each image was counted for the total number of cysts (A, using the phase-brightfield channel) and the number of red cysts (B, using the TRITC channel). The percent of total cells staining red with EthD-1 was then calculated (C). Chlorhexidine was dissolved in DMSO, $H_2O_2$ was dissolved in PBS. Two-fold dilutions were prepared, and drugs were added at 1% final volume in PBS. The mean and SEM of 3 replicates per condition are shown.

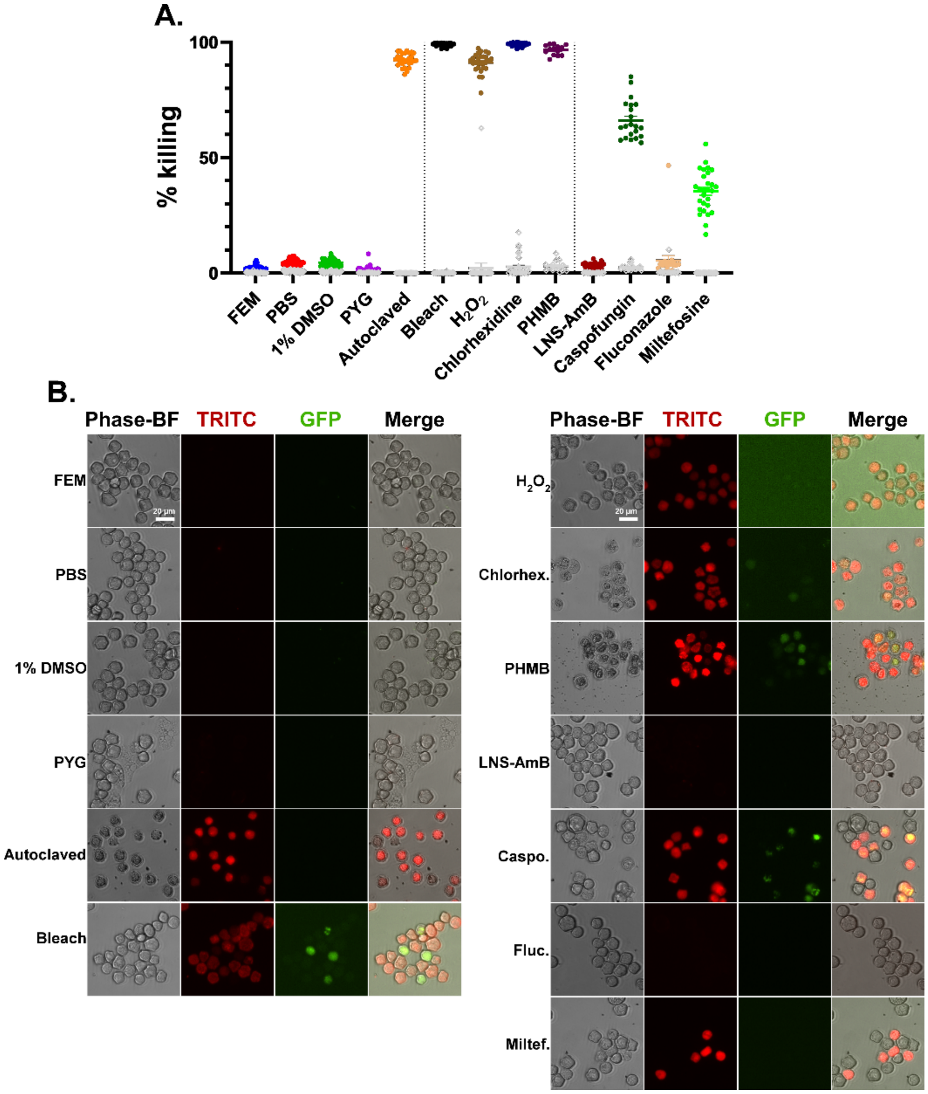


**Figure S7: EthD-1 accurately stains dead cysts while c-AM labels very few cysts. A:** Scatter plot (colored points) reports percent killing (100% - %survival) for the same data shown in figure 3, measured by the percent of EthD-1-stained cysts. Grey diamonds represent the percent of cells dually labeled with red and green fluorescent stains for each condition. The mean and SEM of at least 3 independent experiments with 7 replicates per condition are shown. **B:** Microscopy of cysts after 20-hour incubation with drugs or media and live/dead staining, as described in figure 3. Images acquired using a Nikon Eclipse Ti-E inverted microscope (20x objective). Representative images displayed.

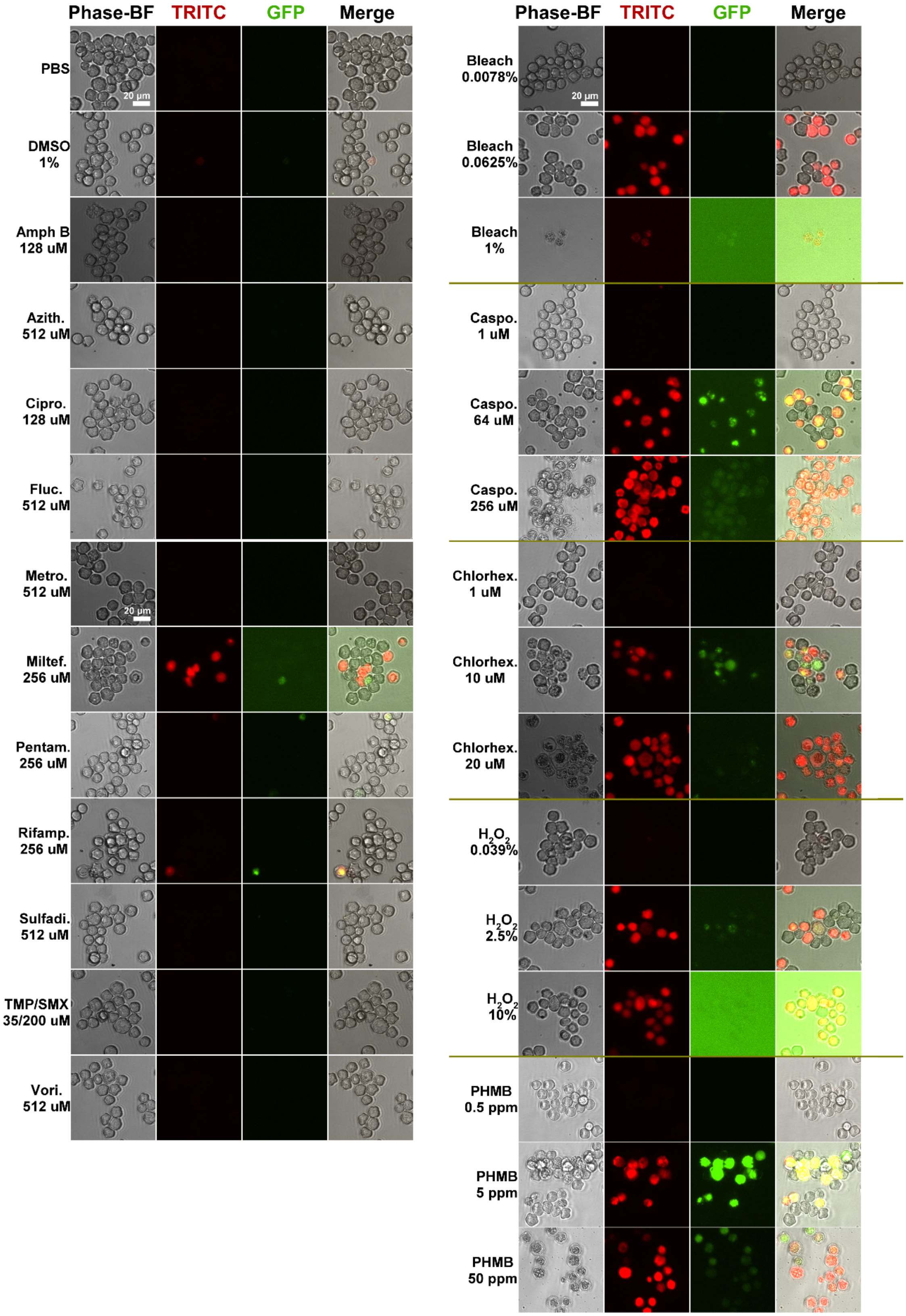
Phase-BF
TRITC
GFP
Merge
PBS
20 μm
DMSO 1%
Amph B 128 uM
Azith. 512 uM
Cipro. 128 uM
Fluc. 512 uM
Metro. 512 uM
20 μm
Miltef. 256 uM
Pentam. 256 uM
Rifamp. 256 uM
Sulfadi. 512 uM
TMP/SMX 35/200 uM
Vori. 512 uM
Phase-BF
TRITC
GFP
Merge
Bleach 0.0078%
20 μm
Bleach 0.0625%
Bleach 1%
Caspo. 1 uM
Caspo. 64 uM
Caspo. 256 uM
Chlorhex. 1 uM
Chlorhex. 10 uM
Chlorhex. 20 uM
H2O2 0.039%
H2O2 2.5%
H2O2 10%
PHMB 0.5 ppm
PHMB 5 ppm
PHMB 50 ppm

**Figure S8: Microscopy corroborates drug testing results.** Microscopy of cysts after 20-hour incubation with drugs or media and live/dead staining, as described in figure 4. Left panel, inactive drugs (highest concentration only shown); right panel, cysticidal agents (lowest, highest, and closest to IC50 concentrations displayed). Images acquired using a Nikon Eclipse Ti-E inverted microscope (20x objective). Representative images displayed.

**Supplemental Tables**

**Table S1: Ingredients tested for new encystment medium.**

| Category | Media component | Concentrations tested | Purpose | References |
|---|---|---|---|---|
| Metals | $CaCl_2$ | 100 µM, 200 µM, 250 µM, 500 µM, 12.5 mM, 50 mM | Cofactor of proteases involved in encystment | 24, 25 |
| | $CoCl_2$ | 100 µM | Cofactor of proteases involved in encystment | 25 |
| | $CoSO_4$ | 100 µM | Cofactor of proteases involved in encystment | 25 |
| | $MgCl_2$ | 100 µM, 10 mM, 50 mM, 100 mM | Cofactor of proteases involved in encystment | 24, 25, 29 |
| | $MgSO_4$ | 50 mM | Cofactor of proteases involved in encystment | 24 |
| | $MnCl_2$ | 100 µM | Cofactor of cellulose synthase | 27 |
| | $MnSO_4$ | 100 µM, 200 µM, 250 µM, 500 µM, 1 mM, 20 mM | Cofactor of proteases involved in encystment, cofactor of cellulose synthase | 25, 27 |
| | $NiCl_2$ | 50 µM, 100 µM, 1 mM | Cofactor of proteases involved in encystment | 25 |
| | $ZnCl_2$ | 100 µM | Cofactor of proteases involved in encystment, cofactor of cellulose synthase | 24, 25, 26 |
| Sugars | Galactose | 7.5 mM, 15 mM, 50 mM | Cyst wall component | 12 |
| | Glucose | 5 mM, 10 mM, 25 mM, 50 mM, 100 mM, 200 mM | Substrate for cellulose synthesis, increase osmolarity, cyst wall component | 12, 28, 29 |
| | Mannose | 1 mM, 10 mM, 17.5 mM, 35 mM, 40 mM | Cyst wall component | 12 |
| Peptides | Casamino Acids (CA) | 0.5%, 1% | Substrate for ectocyst protein synthesis | 9 |
| | Hydroxy-proline (OH-pro) | 1 mM, 10 mM | Cyst wall component | 12 |
| Other | ATP | 10 µM, 100 µM | Galactokinase substrate, activate glycogen phosphorylase (through cAMP) | 12, 22 |
| | DMSO | 0.5% | Stress | 22 |
| | $H_2O_2$ | 0.005% | Stress | 22 |
| | NaCl | 25 mM, 50 mM | Increase osmolarity | 28, 33 |

| | | | | |
|---|---|---|---|---|
| | NAD | 100 µM, 1 mM, 10 mM | Cofactor of sirtuin involved in encystment | 23 |
| | PEG 3000 | 1%, 5% | Increase osmolarity | 28, 33 |

**Table S2: Top 45 encystment medium formulations.**

| Media component | $CaCl_2$ (100 μM) | $MgCl_2$ (50 mM) | $MnSO_4$ (100 μM) | $NiCl_2$ (50 μM) | Glucose (50 mM) | Mannose (35 mM) | CA (0.5%) | OH-pro (10 mM) | NAD (100 μM) |
|---|---|---|---|---|---|---|---|---|---|
| Media # | | | | | | | | | |
| 1 | | | | | x | x | x | | |
| 2 | x | | | | x | x | x | | |
| 3 | 200 μM | | | | x | x | x | | |
| 4 | 200 μM | x | | | | x | | | |
| 5 | | x | 200 μM | | | x | | x | |
| 6 | 200 μM | x | | | | x | | x | |
| 7 | 200 μM | x | 200 μM | | | x | | x | |
| 8 | 200 μM | x | | | x | x | | | |
| 9 | 200 μM | x | 200 μM | | x | x | | | |
| 10 | | x | 200 μM | | x | x | | x | |
| 11 | 200 μM | x | | | x | x | | x | |
| 12 | 200 μM | x | 200 μM | | x | x | | x | |
| 13 | | x | | | x | | | | |
| 14 | x | x | | | x | | | | |
| 15 | | x | | | x | | | x | |
| 16 | | x | x | | x | | | x | |
| 17 | x | x | | | x | | | x | |
| 18 | x | x | x | | x | | | x | |
| 19 | | x | | | | x | | | |
| 20 | x | x | | | | x | | | |
| 21 | | x | | | | x | | x | |
| 22 | | x | x | | | x | | x | |
| 23 | x | x | | | | x | | x | |
| 24 | x | x | x | | | x | | x | |
| 25 | | x | | | x | x | | | |
| 26 | | x | x | | x | x | | | |
| 27 | x | x | | | x | x | | | |
| 28 | x | x | x | | x | x | | | |

| | | | | | | | | | |
|---|---|---|---|---|---|---|---|---|---|
| 29 | | x | | | x | x | | x | |
| 30 | x | x | | | x | x | | x | |
| 31 | x | x | x | | x | x | | x | |
| 32 | x | | | x | x | | | x | |
| 33 | | x | x | x | x | | | x | |
| 34 | x | x | | x | x | | | x | |
| 35 | | x | | x | | x | | x | |
| 36 | | | | | | | | x | |
| 37 | | | x | | | | | x | |
| 38 | x | | | | | | | x | |
| 39 | x | x | x | | | | | x | |
| 40 | | x | | x | | | | | |
| 41 | x | x | | x | | | | | |
| 42 | | x | | | | | | x | x |
| 43 | | x | x | | | | | x | x |
| 44 | x | x | | | | | | x | x |
| 45 | x | x | x | | | | | x | x |

Listed ingredients were added to the base medium EMb for all formulations. Media #1-35 (corresponding with the numbered conditions in figure S4) encysted for 72 hours in 150 µL per well; media #36-45 encysted for 48 hours in 100 µL per well. Concentrations tested are listed at the top of each column unless otherwise indicated. OH-pro, hydroxyproline; CA, casamino acids.

### <u>Supplemental methods</u>

#### Media

10X PBS: 80 g/L NaCl, 2.0 g/L KCl, 14.4 g/L $Na_2HPO_4$, 2.4 g/L $KH_2HPO_4$ in distilled $H_2O$, diluted to 1X working concentration by diluting in distilled $H_2O$.

Flynn's Encystment Medium (FEM): EMb supplemented with 50 mM glucose, 10 mM hydroxyproline, 100 µM $MnSO_4$, 50 µM $NiCl_2$, 50 mM $MgCl_2$ in distilled $H_2O$; stored in plastic bottles at 4˚C, warmed to room temperature before adding to cells. Protocol available at https://dx.doi.org/10.17504/protocols.io.kxygxqx14v8j/v1.

All other media as described in (5).

All media were sterilized by passage through a 0.22 µm filter except for LB, which was autoclaved.

#### SRB assay

Plates fixed with 10% TCA were incubated at 4˚C for at least one hour, washed four times by submerging in tap water, tapped dry on paper towels, and air dried. Cells were stained with 50 µL per well 4% (w/v) SRB dye in 1% acetic acid, incubated in the dark for 15 minutes at room temperature, and washed of excess dye by submerging three times in 1% acetic acid. Plates were tapped dry on paper towels and air dried. Bound SRB dye was solubilized by adding 150 μL per well 10 mM Tris-HCl (pH 8.0), incubating on gyrating platform for 5 minutes at room temperature, and measuring the absorbance in a Tecan Infinite M200 Pro plate reader with the following program: linear shake (1 mm amplitude) 30 seconds, wait 5 seconds, measure absorbance at 565 nm (25 flashes/read) (protocol at dx.doi.org/10.17504/protocols.io.bvpen5je).
